# NanoBlot: A Simple Tool for Visualization of RNA Isoform Usage From Third Generation RNA-sequencing Data

**DOI:** 10.1101/2022.10.26.513894

**Authors:** Samuel DeMario, Kevin Xu, Kevin He, Guillaume Chanfreau

## Abstract

RT-PCR and Northern blots have long been used to study RNA isoforms usage for single genes. Recently, advancements in long read sequencing have yielded unprecedented information about the usage and abundance of these RNA isoforms. However, visualization of long-read sequencing data remains challenging due to the high information density. To alleviate these issues we have developed NanoBlot, a simple, open-source, command line tool, which generates Northern blot and RT-PCR-like images from third generation sequencing data. NanoBlot accepts processed bam files. Plotting is based around ggplot2 and is easily customizable. Advantages of NanoBlots include: designing probes to visualize isoforms which would be impossible with traditional RT-PCR or Northern blots, excluding reads from the Nanoblots based on the presence or absence of a specified region and, multiplexing plots with multiple colors. We present examples of NanoBlots compared to actual northern blot data. In addition to traditional gel-like images, NanoBlot also outputs other visualizations such as violin plots. The use of Nanoblot should provide a simple answer to the challenge of visualization of long-read RNA sequencing data.

## Introduction

mRNA isoforms usage is biologically important and dynamic. Alternate splicing, alternate polyadenylation site selection and alternative transcription start site selection, all result in the production of different mRNA isoforms from the same transcription unit. These diverse isoforms increase proteome complexity and the presence of regulatory elements in each isoform can impact steady-state expression levels. RNA isoforms usage has long been studied on the scale of individual genes, first predominantly with Northern blots and more recently using RT-PCR. While easily performed, RT-PCR and Northern blots have limitations. RT-PCR cannot distinguish between isoforms with differences outside of the region targeted for PCR. Northern blots detect different isoforms which hybridize to the same probe, but they can only detect one or two gene products at a time. In addition, Northern blots cannot easily visualize isoforms that differ by only a few nucleotides if the target RNA is long. The development of long-read sequencing technologies is an exciting innovation which allows for high throughput studies of isoform usage.

However, due to the information density, visualization of long-read sequencing data remains challenging and tools for visualization are scarce. One common approach to visualization is using images from genome browsers such as the Integrative Genome Viewer [1] or UCSC Genome Browser (http://genome.ucsc.edu) [2]. This has the advantage of showing all the reads obtained in a specific genomic region. However, for genes with many isoforms or long introns relative to coding regions compact visualization is challenging. Tracks can be edited to shorten long introns and reads can be down sampled to increase information density. Tools such as ScisorWiz [3] help by automating much of this process. However, each additional sample requires a new track to be added. This takes up considerable space and makes representation of more than a few samples in a single image non-feasible. Another common approach is assigning each transcript to a discrete isoform. Tools such as IsoTV [4] and Swan [5] toolkit automate this process and help create intuitive visualizations of isoforms. However, this requires detailed annotation of the host genome and assigns reads to distinct groups. This means that if the model organism being investigated is not well studied, or if there are continuous variations in isoform lengths, then isoform counting is not possible. A possible solution to these limitations is to represent each read as a datapoint representing the length of that read. The result of this is a plot which looks similar to an RT-PCR gel or a Northern blot [6]. To explore this idea, we have developed NanoBlot, a simple command line tool which allows for easy visualization of long read sequencing data in an intuitive form reminiscent of Northern blots or RT-PCR data. The most recent version of NanoBlot can be found on GitHub (https://github.com/SamDeMario-lab/Nano_blot).

## Results and Discussion

### Main Features of NanoBlot

NanoBlot was originally developed for visualization of RNA isoforms detected using Oxford Nanopore (ONT) long read sequencing datasets. However, NanoBlot should be able to handle data obtained with other long read sequencing techniques such as PacBio. NanoBlot takes aligned, positionally-sorted, and indexed BAM files as input (Figure 1A). The location of the input data should be provided via a tab-separated metadata table. The table must include a unique name for each sample and the location of the aligned BAM files. An example metadata table is provided in the supplementary materials. The sequencing data used does not need to be normalized prior to running NanoBlot as normalization is included. NanoBlot requires a series of target genomic regions referred to as “probes” (Figure 1A). Probes are supplied as a standard six column BED file. Finally, NanoBlot requires a plot metadata table listing: A unique name for the resulting plot, the BAM files in the order in which they should be loaded, one or more probes to be used, and an optional field which specifies any regions which should be excluded (referred to as “antiprobes”). NanoBlot then subsets each of the input bam files to only the sequencing reads which overlap with the specified probe region (Figure 1A) [7]. If multiple probes are specified, then each read must overlap with all specified regions to be included as AND statements (see README for further information). If antiprobes are included, NanoBlot removes any reads which map to the antiprobe(s) region. This is an optional function which can be used to exclude specific reads as a means to simplify plots. Finally, NanoBlot outputs the raw data and generates three different plots: (i) A plot where each read is represented as a line corresponding to its length. This type of plot is the most reminiscent of an actual Northern blot or RT-PCR gel (Figure 1B); (ii) a violin plot which is useful for representing samples where the isoforms of interest have significantly different sizes (Figure 1B); or (iii) a ridge plot where density plots for each sample are slightly overlapped, making it useful for showing subtle differences in length (Figure 1B). NanoBlot can also accept user generated R scripts for additional customization.

**Figure 1.**
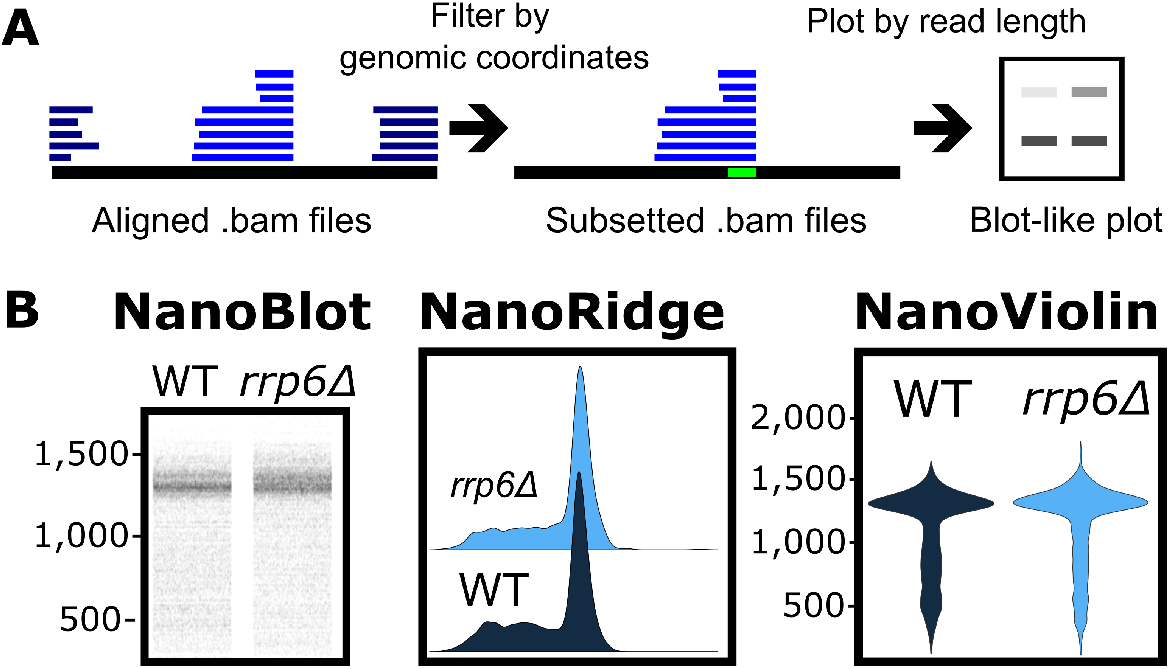
Overview of NanoBlot and Example Plots. A) General overview of NanoBlot. NanoBlot’s first step is subsetting of BAM files based on a genomic region, shown in bright green. Reads are then represented as bands based on the length of the reads. Multiple samples and probes can be show on the same plot. B) Three types of plots defaultly produced by NanoBlot.

### Control of Plotting and features for Northern blot like Figures

Typical blot-like pictures were generated using NanoBlot and compared to actual northern blot data for RNAs extracted from Saccharomyces cerevisiae wild-type and rrp6Δ mutant (Figures 2A, 2B). The pictures generated by NanoBlot were strikingly similar to actual Northern blot data, even for low abundance products such as an RNA cleavage product present for *RPL18A* in the *rrp6*Δ mutant (Figure 2A), or for low abundance isoforms of the *ADI1* transcripts (Figure 2B). In a NanoBlot the scale of the y-axis representing the size of the RNA isoforms can be changed allowing for more precise control over the distribution of bands. By default, NanoBlots are scaled to include the full range of lengths, however, this occasionally results in plots that appear squished. Setting a custom length range can also increase plot legibility. If isoforms with large size differences are to be represented on the same blot, a logarithmic scale can be used for the y-axis (Figure 2B). Traditional Northern blots typically rely on the use of a single radioactive or fluorescent probe which detects RNAs hybridizing to the probe. Membrane stripping and subsequent hybridization can be performed to detect different RNAs (e.g., a loading control). By contrast, NanoBlots can be overlapped and represented in multiple colors making multiplexing possible. To illustrate this feature, we used data from wild-type and *rrp6*Δ mutant to generate blot like pictures representing the relative abundances of the *RPS7B* and *snR4* RNAs in these strains (Figure 2C). NanoBlot can theoretically be multiplexed infinitely. However, overplotting quickly becomes a concern. This can be partially alleviated by separating the probes into adjacent lanes, as shown for *RPS7B* and *snR4* in Figure 2D. Bands in NanoBlots are not at risk of being masked by nearby highly abundant isoforms allowing for visualization of isoforms with similar sizes. However, it is worth noting that NanoBlots do not give single nucleotide resolution. Limitations in base calling and mapping efficiency can also result in small discrepancies in transcript length. For instance, the mature form of *snR37* is 386 nts long however, Nanopore data shows a band slightly below the expected size (Figure 2E).

**Figure 2.**
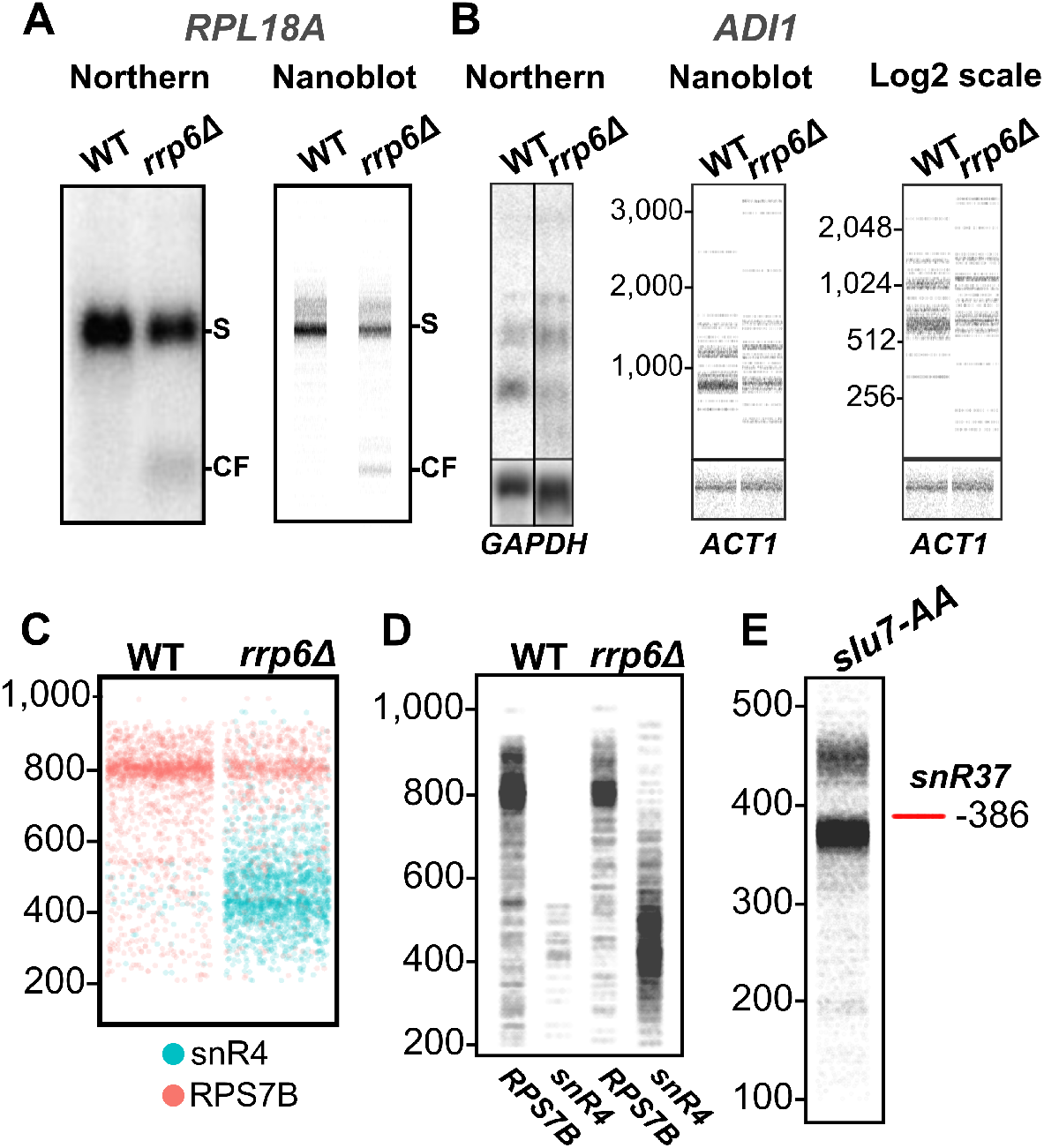
Examples of NanoBlots, Comparison to Northern blots, Multiplexing, and Size Estimation. A) Comparison between 32P Northern Blot and NanoBlot targeting the 5’ Exon of *RPL18A*. Notice the accumulation of the small molecular weight species in the rrp6d samples. S:Spliced *RPL18A* mRNA, CF:5’ cleave fragment B) Example NanoBlot on the low abundance gene, *ADI1*. Note the strong banding pattern. Also shown is a NanoBlot with the size axis shown on a log2 scale. C) Multiplex NanoBlot showing 2 genes probed with different colors. D) The same data as in C but with each probe + samples combination separated into its own lane. E) NanoBlot showing size misrepresentation of *snR37*. The red line indicates the true size of the mature snoRNA. *slu7-AA*: Direct RNA sequencing of *Slu7-FRB* rapamycin treated RNAs pre-processed with Terminator exonuclease, rSAP, and in-vitro poly-adenylated.

### Conditional Probes

One of the exciting possibilities of NanoBlots is that NanoProbes can be designed to select for more specific targets than what is possible with Northern blots or RT-PCR. NanoProbes can be designed to ‘‘hybridize” with reads in a conditional manner. As an example, one may wish to investigate snoRNA processing. A probe can be designed to hybridize with a specific read only if it contains the mature snoRNA sequence AND has a 3’ extension, excluding mature snoRNAs lacking a 3’ extension. This feature is impossible to replicate using traditional Northern blots. NanoProbes can also be designed to exclude reads which map to a particular region. For example, if one wanted to look at the isoform distribution of a sub-population of a gene which does not have a specific exon, a probe could be designed to select for this.

### Quantification of Extended isoforms

NanoBlots are particularly suited to representing isoforms without discrete lengths. We illustrate this feature by highlighting the accumulation of polyadenylated extended forms of unprocessed or partially processed forms of snoRNAs in the *rrp6*Δ mutant. The 3’ ends of many pre-snoRNAs are generated by the NNS machinery in a stochastic manner. The extended species are then processed by the nuclear exosome with the assistance of the TRAMP complex [8]. The deletion of *RRP6* results in the general accumulation of polyadenylated, extended species of *snR37* whereas the deletion of *TRF4* results in the accumulation of a discrete larger extended species. In a traditional Northern blot, the similar sizes of these two discrete products would result in the bands appearing to merge, however in a NanoRidge plot it becomes clear that there is a bimodal distribution (Figure 3A).

**Figure 3.**
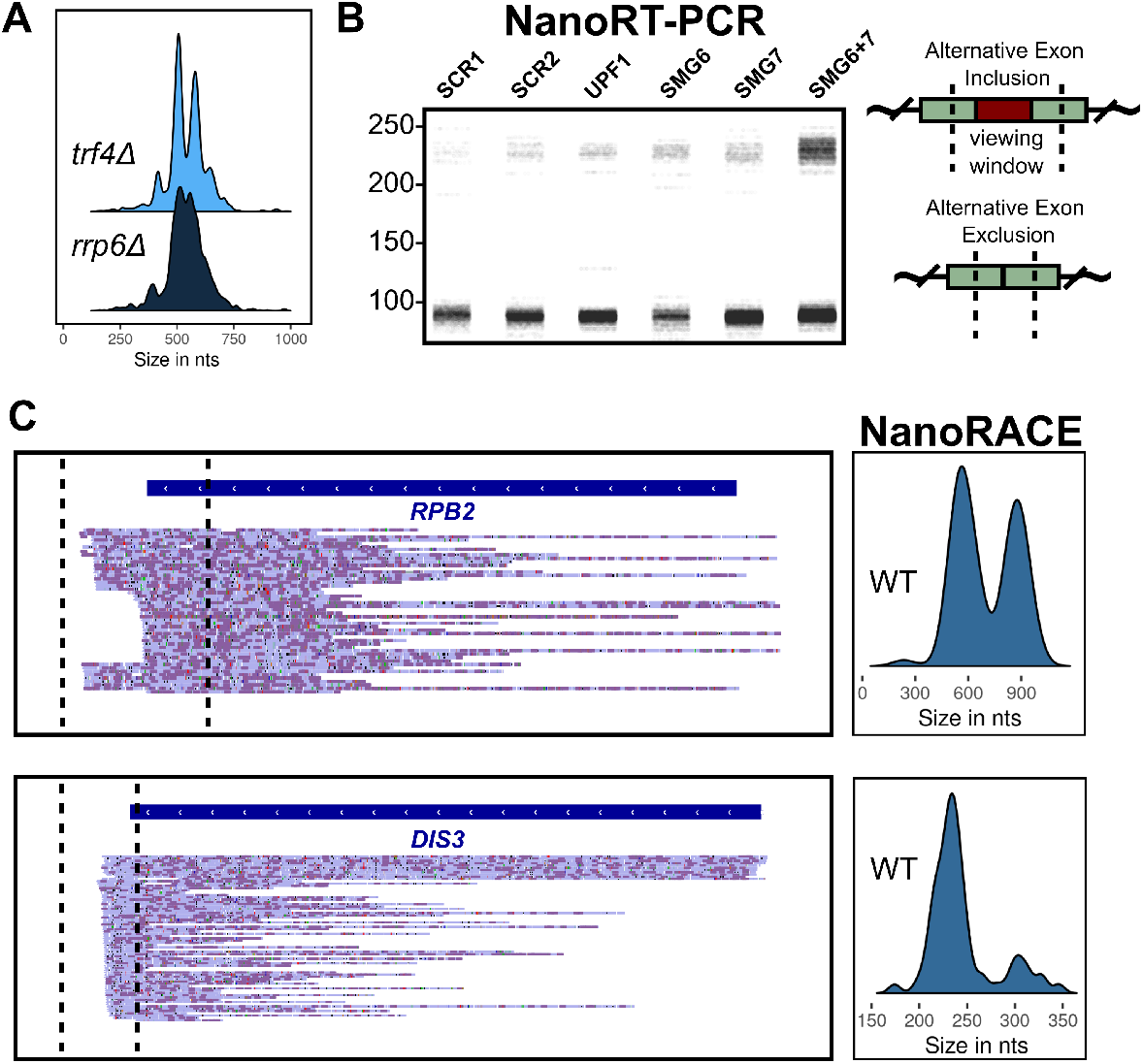
NanoRidge, NanoRT-PCR and, NanoRACE. A) NanoRidge showing a clear increase in bimodality in the *trf4*Δ samples verses the *rrp6*Δ samples. B) NanoRT-PCR plot showing an alternative exon inclusion event. SCR1: Scramble 1, SCR2: Scramble 2, UPF1: UPF1 Knockdown, SMG6: SMG6 Knockdown, SMG7: SMG7 Knockdown, SMG6+7: SMG6 and SMG7 Double Knockdown. Diagram on the right showing the approximate position of the viewing window around the alternative exon (Not shown to scale). C) IGV screenshots of WT sequencing data and NanoRACE plots. *RPB2* shows alternative polyadenylation site usage while *DIS3* shows a single polyA site. Note the abundance of short reads making representation via a NanoBlot challenging.

### NanoRT-PCR and NanoRACE

NanoBlot can also produce RT-PCR like plots. To produce an RT-PCR plot a viewing window needs to be specified for each plot. This is done by inputting a single BED file entry in the plot-metadata file for each RT-PCR NanoBlot. NanoBlot then checks each read to ensure that they overlap with the beginning AND end of the viewing window. Any read which does not is excluded from the plot. Finally, NanoBlot counts the mapped bases inside of the viewing window for each read and plots them as normal. To illustrate this feature, we generated an RT-PCR like plot to visualize the accumulation of an isoform containing an alternate exon inclusion event for the *BAG1* gene. This isoform contains a premature translation termination codon and can be detected upon siRNA knockdowns of human nonsense-mediate mRNA decay (NMD) factors [9], particularly when the NMD factors *SMG6* and *SMG7* are co-depleted (Figure 3B). NanoRACE is a slight variation on NanoRT-PCR. The difference is that NanoRACE does not remove reads without bases mapping to the 3’ end of the viewing window while NanoRT-PCR does. This is a useful tool for studying alternative polyadenylation site usage. Most of the reads for *RPB2* and *DIS3* are not full length meaning they cannot be used for accurate size estimation. However, they still have accurate information about poly(A) site usage. By setting the viewing window of a NanoRACE plot to cover the poly(A) sites accurate density plots are produced. *RPB2* has 2 poly(A) sites used at approximately equal rates while DIS3 has only a single poly(A) site (Figure 3C).

## Advantages and Limitations of NanoBlots

### Issues linked to cross-hybridization

One advantage of NanoBlot is that cross-hybridization issues are alleviated. Fundamentally, cross-hybridization issues in Northern blots are due to similarities between the probe sequence and other RNA species present in the sample. The presence of ribosomal RNAs in northern blots performed from total RNAs tend to generate cross-hybridization signals due to the high abundance of these RNAs. In standard Northern blots, cross-hybridization issues can be alleviated by using probes with a higher degree of specificity to the target region and/or by using more stringent hybridization and/or washing conditions after hybridization. However, it is not always possible to implement these solutions, especially if the target sequence is limited in length. One example would be the detection of the splice isoforms that are expressed for a gene which contain a specific, short exon. One could design a probe that targets the specific exon, but the maximum length of the probe is fundamentally limited by the length of the target exon, which can be problematic for microexons or some cassette exons that have very slight sequence differences. In a NanoBlot the entire length of the transcript can be used to map to the genome and then the data can be filtered based on the presence of a specific region.

### Issues and limitations linked to repeated or homologous sequences

NanoBlots are not without limitations. Reads can be mismapped or rendered unmappable. Of particular concern is genes with multiple copies or paralogues. As an example, the *S.cerevisiae TDH1* gene has 2 closely related paralogues, *TDH2* and *TDH3*. Third generation aligners (e.g., Minimap2) have a particularly difficult time uniquely mapping these reads and therefore they are either not represented in the results BAM files or flagged as non-uniquely mapped. As a result, NanoProbes targeting *TDH1* can result in blank plots depending on how the mapping was handled.

### Sensitivity

NanoBlots are subject to all the same limitations of Nanopore sequencing. One of the major issues is that short RNA fragments or isoforms are not efficiently mapped. As a result, smaller fragments cannot be efficiently represented in a NanoBlot. Sensitivity is also a concern. It is challenging to quantify how sensitive Nanopore sequencing is compared to Northern blots. As an example, *ADI1*, a low abundance RNA transcript, is shown as either a 32P Northern blot, or a NanoBlot (Figure 2B). As with any library prep there is potential biases introduced by the ONT sequencing kits. As new strategies are developed to combat biases, they can be seamlessly incorporated into NanoBlot.

### Issues linked to the presence of degradation products

RNA degradation and incomplete transcript sequencing are a problem for accurate transcript size estimation. Transcripts partially degraded from the 5’ end but with an intact poly(A) tail will still be sequenced via ONT sequencing resulting in shortened read sequences. In NanoBlots, this results in a low molecular weight smear which can make representation difficult especially for longer transcripts. One possible solution is to specify 2 regions as probes, one at the 5’ end of the transcript and another at the 3’ end. This will result in the visualization of only near full length transcripts. If isoforms of substantially different lengths are being investigated this type of selection could result in the blot being biased towards shorter transcripts.

### Detection and Visualization of polyadenylated and non-polyadenylated species and poly(A) tail length

Standard Nanopore RNA sequencing is poly(A) based meaning that any transcript lacking a poly(A) tail will not be sequenced. There do exist ways to prepare RNAs for sequencing non-poly(A) RNAs. RNAs can be tailed or in-vitro polyadenylated prior to library synthesis[10, 11]. However, some biologically relevant RNAs lack 3’ hydroxyls (e.g., snoRNAs, tRNAs). Treatment with a phosphatase such as rSAP or AnP can allow these RNAs to be sequenced. However, if enzymatic treatments are preformed samples can no longer be used quantitatively because of possible biases caused by the enzymatic treatments. Poly(A) tail length is not included in the bands shown on NanoBlots. The poly(A) tail length could theoretically be included if a direct RNA sequencing strategy is used, because poly(A) tail length can be estimated and added to the length although the initial release of NanoBlot does not have this functionality[12].

## Methods

### Yeast growth

Wild-type and *RRP6* knockout yeast strains are from the BY4741 genetic background. Yeast cultures were grown in YPD (1% w/v yeast extract, 2% w/v peptone, and 2% w/v dextrose). Briefly, 50ml cultures were grown at the standard 30°C to an OD600 of 0.4. Cells were then pelleted and flash frozen in liquid nitrogen for RNA isolation. RNA isolation was performed as described in Wang et al[13].

### Library prep

Total RNAs were treated with DNAse I (Invitrogen, catalog #: 18-068-015) following the manufacturer’s protocol. Using the direct RNA sequencing kit from Oxford Nanopore (ONT, catalog #: SQK-RNA002) libraries were prepared according to the manufacturer’s instructions. Sequencing was performed using R9.4 flow cells on a MinION Mk1B device and sequenced for 48 hours. Basecalling was performed using Guppy Basecaller (Version 6.1.1+1f6bfa7f8). Reads were then mapped to the Saccharomyces cerevisiae genome: (S288C_reference_sequence_R64-3-1) using Minimap 2 (Version 2.17-r941).

### Normalization

Because NanoBlots are generated from full RNA sequencing datasets many different normalization techniques are viable. By default NanoBlot accepts an annotation file as input and normalization is done via DESeq2[14]. First NanoBlot generates counts tables for each input file using HTSeq[15]. Next NanoBlot calculates normalization factors via DESeq2. Finally, NanoBlot generates plots with increased sampling for smaller libraries. Unnormalized data is used to produce density plots. If an annotation file is not available, NanoBlot can also generate normalization factors based solely on library depth. Other normalization techniques can be used depending on the specific datasets used. NanoBlot can accept either normalized or unnormalized reads.

### Dataset Generation

Bedtool’s intersect function is used to subset the input files to only reads which overlap with the region of interest. An arbitrary number of probes can be specified and will be treated as AND probes, requiring each read to overlap with all specified probes. Probes to be used for NOT statements are placed in the optional ‘antiprobe’ column of the plot_metadata file. OR logical statements must be done as two separate selections and be merged by the end user. NanoBlot can also produce RT-PCR or RACE like plots where the length of each read is counted as the number of mapped bases in a viewing window. The ‘-Y’ flag is used to specify the RT-PCR function. It is important to note that the plot_metadata file for RT-PCR/RACE and standard NanoBlot usage are slightly different, see README.md for additional information. At the time of publication there is a known bug where for long genomes NanoRT-PCR and NanoRACE do not function properly, see the GitHub repo for the most up to date information.

### R and Plotting

All required R packages are installed via Bioconductor[16]. The subsetted BAMs are read into R using scanBAM() from the Rsamtools package[17]. The length of each read is calculated from the CIGAR string and listed in the qwidth column. Data is reformatted for plotting with the dplyr[18] package. Plot generation is done via ggplot2[19, 20]. Custom R scripts can be passed to NanoBlot for further customization. The NanoBlot script accepts the ‘-F’ flag which skips resubsetting of bam files if only plotting is required. The default R script also writes out a .RDS file with the data used to generate each plot, this can be easily read into an Rscript for testing.

## Data Availability

Data for Saccharomyces cerevisiae *trf4*Δ was downloaded from the ENA, ascension number: SAMEA7708082. Data for Saccharomyces cerevisiae *SLU7* Anchor-Away was downloaded from the SRA, ascension number: PRJNA827814. Data for human NMD knockdown was downloaded from the ArrayExpress database at EMBL-EBI (www.ebi.ac.uk/arrayexpress) under accession number: E-MTAB-10452. All custom R script used to generate figures in this publication are available in the NanoBlot GitHub (https://github.com/SamDeMario-lab/Nano_blot). Sequencing data generated for this study are available at the SRA, ascension number: PRJNA891745. NanoBlot is available under the MIT license. DOI: 10.5281/zenodo.7213547

## Competing interests

The authors declare that they have no competing interests.

## Author’s contributions

SD and KX authored the code. SD and GC conceived the study and authored the manuscript. KH generated sequencing libraries and preformed beta testing. SD created figures. All authors reviewed the final manuscript.

## Funding

This work was supported by NIGMS grant GM130370 to GC. KX was supported by the James and Meredith Henry Estate Endowment.

## Notes

https://github.com/SamDeMario-lab/Nano_blot

### Competing Interest Statement

The authors have declared no competing interest.

## References

[1] James T Robinson et al. “Integrative genomics viewer”. In: Nature Biotechnology 29.1 (Jan. 1, 2011), pp. 24–26. ISSN: 1546-1696. DOI: 10.1038/nbt.1754. URL: https://doi.org/10.1038/nbt.1754.

[2] W. James Kent et al. “The human genome browser at UCSC.” In: Genome research 12.6 (June 2002), pp. 996–1006. ISSN: 1088-9051. DOI: 10.1101/gr.229102.

[3] Alexander N Stein et al. “ScisorWiz: visualizing differential isoform expression in single-cell long-read data”. In: Bioinformatics 38.13 (July 1, 2022), pp. 3474–3476. ISSN: 1367-4803. DOI:10.1093/bioinformatics/btac340. URL: https://doi.org/10.1093/bioinformatics/btac340 (visited on 09/12/2022).

[4] Siddharth Annaldasula, Martyna Gajos, and Andreas Mayer. “IsoTV: processing and visualizing functional features of translated transcript isoforms”. In: Bioinformatics 37.18 (Sept. 15, 2021), pp. 3070–3072. ISSN: 1367-4803. DOI: 10.1093/bioinformatics/btab103. URL: https://doi.org/10.1093/bioinformatics/btab103 (visited on 09/12/2022).

[5] Fairlie Reese and Ali Mortazavi. “Swan: a library for the analysis and visualization of long-read transcriptomes”. In: Bioinformatics 37.9 (May 1, 2021), pp. 1322–1323. ISSN: 1367-4803. DOI: 10.1093/bioinformatics/btaa836. URL: https://doi.org/10.1093/bioinformatics/btaa836 (visited on 09/12/2022).

[6] ArnaudAU Guilcher et al. “Full Length Transcriptome Highlights the Coordination of Plastid Transcript Processing”. In: International Journal of Molecular Sciences 22.20 (2021). ISSN: 1422-0067. DOI: 10.3390/ijms222011297.

[7] Aaron R. Quinlan and Ira M. Hall. “BEDTools: a flexible suite of utilities for comparing genomic features.” In: Bioinformatics (Oxford, England) 26.6 (Mar. 15, 2010), pp. 841–842. ISSN: 1367-4811 1367-4803. DOI: 10.1093/bioinformatics/btq033.

[8] Pawel Grzechnik and Joanna Kufel. “Polyadenylation Linked to Transcription Termination Directs the Processing of snoRNA Precursors in Yeast”. In: Molecular Cell 32.2 (Oct. 24, 2008), pp. 247–258. ISSN: 1097-2765. DOI: 10.1016/j.molcel.2008.10.003. URL: https://www.sciencedirect.com/science/article/pii/S1097276508006850.

[9] Evangelos D. Karousis et al. “Nanopore sequencing reveals endogenous NMD-targeted isoforms in human cells”. In: Genome Biology 22.1 (Aug. 13, 2021), p. 223. ISSN: 1474-760X. DOI: 10.1186/s13059-021-02439-3. URL: https://doi.org/10.1186/s13059-021-02439-3.

[10] Yanru Liu et al. “Splicing inactivation generates hybrid mRNA-snoRNA transcripts targeted by cytoplasmic RNA decay”. In: Proceedings of the National Academy of Sciences 119.31 (Aug. 2, 2022). Publisher: Proceedings of the National Academy of Sciences, e2202473119. DOI: 10.1073/pnas.2202473119. URL: https://doi.org/10.1073/pnas.2202473119 (visited on 10/12/2022).

[11] Agnieszka Tudek et al. “Global view on the metabolism of RNA poly(A) tails in yeast Saccharomyces cerevisiae”. In: Nature Communications 12.1 (Aug. 16, 2021), p. 4951. ISSN: 2041-1723. DOI: 10.1038/s41467-021-25251-w. URL: https://doi.org/10.1038/s41467-021-25251-w.

[12] Maximilian Krause et al. “tailfindr: alignment-free poly(A) length measurement for Oxford Nanopore RNA and DNA sequencing.” In: RNA (New York, N.Y.) 25.10 (Oct. 2019), pp. 1229–1241. ISSN: 1469-9001 1355-8382. DOI: 10.1261/rna.071332.119.

[13] Charles Wang et al. “Rrp6 Moonlights in an RNA Exosome-Independent Manner to Promote Cell Survival and Gene Expression during Stress.” In: Cell reports 31.10 (June 9, 2020), p. 107754. ISSN: 2211-1247. DOI: 10.1016/j.celrep.2020.107754.

[14] Michael I. Love, Wolfgang Huber, and Simon Anders. “Moderated estimation of fold change and dispersion for RNA-seq data with DESeq2”. In: Genome Biology 15.12 (Dec. 5, 2014), p. 550. ISSN: 1474-760X. DOI: 10.1186/s13059-014-0550-8. URL: https://doi.org/10.1186/s13059-014-0550-8.

[15] Givanna H Putri et al. “Analysing high-throughput sequencing data in Python with HTSeq 2.0”. In: Bioinformatics 38.10 (May 15, 2022), pp. 2943–2945. ISSN: 1367-4803. DOI: 10.1093/bioinformatics/btac166. URL: https://doi.org/10.1093/bioinformatics/btac166 (visited on 09/12/2022).

[16] W. Huber et al. “Orchestrating high-throughput genomic analysis with Bioconductor”. In: Nature Methods 12.2 (2015), pp. 115–121. URL: http://www.nature.com/nmeth/journal/v12/n2/full/nmeth.3252.html.

[17] M Morgan et al. Rsamtools: Binary alignment (BAM), FASTA, variant call (BCF), and tabix file import. Version 2.12.0. URL: https://bioconductor.org/packages/Rsamtools.

[18] Hadley Wickham et al. dplyr: A Grammar of Data Manipulation. 2022. URL: https://CRAN.R-project.org/package=dplyr.

[19] Hadley Wickham. ggplot2: Elegant Graphics for Data Analysis. Springer-Verlag New York, 2016. ISBN: 978-3-319-24277-4. URL: http://ggplot2.org.

[20] Claus O. Wilke. ggridges: Ridgeline Plots in ‘ggplot2’. 2021. URL: https://CRAN.R-project.org/package=ggridges.

